# Generalization asymmetry in multivariate cross-classification: When representation A generalizes better to representation B than B to A

**DOI:** 10.1101/592410

**Authors:** Job van den Hurk, Hans P. Op de Beeck

## Abstract

In recent years, the use of multivariate cross-classification (MVCC) has grown in popularity as a way to test for the consistency of information in neural patterns of activation across cognitive states. In this approach, a classification algorithm is trained on dataset A and then tested on a different dataset B in order to test for commonalities in the information representation in both datasets. Interestingly, several papers report an asymmetry in the generalization direction: training on A and testing on B returns significantly better decoding results than training on B and testing on A. Whereas several neurocognitive hypotheses have been put forward as an explanation for this phenomenon, none of them has been demonstrated directly. Through simple simulations, we show that asymmetry can arise as soon as two datasets with identical ground truths have a different signal-to-noise ratio (SNR) – generalization is best from the lower SNR to the higher SNR dataset. The extent of the asymmetry is further modulated by the overlap in informative voxels and whether the two datasets have an equal number of informative voxels. These findings demonstrate that the observation of decoding direction asymmetry in MVCC can be explained by simple SNR differences and not necessarily implies complex neurocognitive mechanisms.

## Introduction

Due to the highly multivariate nature of functional magnetic resonance imaging (fMRI) datasets, advanced analysis tools that employ the many dimensions of the data grow increasingly more popular among neuroscientists. These so-called multi-voxel pattern analysis (MVPA) methods have proven to be an effective way to discriminate between distributed and/or spatially fine-grained neural responses (Formisano et al., 2008b; Haxby, 2001; Haynes and Rees, 2006; Kamitani and Tong, 2005; Kriegeskorte et al., 2007; Op de Beeck et al., 2010). Binary classification methods such as linear discriminant analysis (LDA) (McLachlan, 1992) and linear support-vector machines (SVM) (Cortes and Vapnik, 1995) can be used to demonstrate that the neural responses associated with two or more stimulus dimensions or mental states are statistically dissimilar.

Typically, each example in the data is expressed as a vector of F features in an F-dimensional space. Each example is labeled as belonging to 1 of a set of experimental conditions or ‘classes’, after which the data is split in a train and test subset. During the subsequent training phase, the training examples belonging to the different classes are mapped onto the multidimensional space, after which the classifier defines an F-1-dimensional hyperplane or separation boundary that optimally separates the data from different classes. The generalizability of this trained model is subsequently quantified by providing the trained model with the test trials, and assessing the ratio of correctly classified examples. A prediction accuracy that is statistically higher than chance indicates that the 2 classes yield meaningful differences in the responses they elicit in the brain region of interest. This process is typically repeated in several cross-validations with different train- and test-subsets, to assess how well the data would generalize to an independent dataset of the same experiment. This procedure allows drawing conclusions on the dissimilarity of neural responses elicited by two experimental classes within the dataset of interest.

Some studies, however, extend the application of MVPA to generalize between two datasets with different conditions. Following Kaplan et al. (2015), we will refer to this procedure as multivariate cross-classification (MVCC). In this approach, one could for example ask if the difference in multi-voxel patterns in the lateral-occipital cortex (LOC) (Malach et al., 1995) between two objects presented visually would generalize to the haptic sense. This question cannot directly be answered by finding discriminatory responses in the haptic modality in LOC: in theory different neural populations could be at work compared to visual processing. However, one could combine both modalities in a single pattern analysis pipeline: the classification algorithm is trained on the neural responses elicited by the visual stimuli, and then asked to predict the presented object in the haptic trials. If this approach yields a successful decoding performance, this constitutes evidence that indeed the same neural patterns in LOC are involved in processing the two object types independent of modality.

Over the last years, a steady increase in studies using this MVCC procedure can be observed in the neuroscience literature (e.g. Awwad Shiekh Hasan et al., 2016; Buchweitz et al., 2012; Cichy and Teng, 2017; Eger et al., 2009; Etzel et al., 2008; Harrison and Tong, 2009; Johnson and Johnson, 2014; Paquette et al., 2018; van den Hurk et al., 2017). Interestingly, several of these studies have reported that the cross-classification performance depends on the decoding direction: training on dataset A and predicting on dataset B can yield significantly different results than vice versa. Here we will refer to this phenomenon as decoding direction asymmetry or DDA.

The proper interpretation and explanation of DDA is unknown. Explanations in the literature range from practical suggestions without much theoretical assumptions up to high-level neurocognitive explanations. An example of the former is the intuitive reasoning of Kaplan et al. (2015) that one direction of generalization might be better than the other because of differences in signal to noise. An example of the latter comes from Etzel et al. (2008) in a study of auditory mirror neurons that involved generalization between auditory and motor stimulation. They suggested that only the subset of neurons with auditory properties would be relevant, which is why they trained on auditory and tested on motor stimulation. Similarly, Eger et al. (2009) observed that the discrimination of numerical magnitudes generalized from training on digits to testing on dots, but not from training on dots to testing on digits. They used the asymmetry as evidence in support of a neural network model of numerical representations that proposed that only a subset of the neurons with selectivity for nonsymbolic magnitudes in dots would have acquired selectivity for symbolic magnitudes. Training on digits would allow the classifier to focus on these neurons/voxels only.

Because the exact interpretation of this generalization asymmetry remains unclear, it is not surprising that the correct way to approach asymmetric results is disputed (Kaplan et al., 2015). Some authors average across decoding directions and report the mean (Man et al., 2012; Oosterhof et al., 2012), others report only one direction (Etzel et al., 2008; Johnson and Johnson, 2014), and some report both directions directly (Akama et al., 2012; Eger et al., 2009; Quadflieg et al., 2011; van den Hurk et al., 2017). Understanding the exact causal factors that lead to DDA can mediate statistical considerations and interpretations of the data.

In the current paper, we use simulations to examine the influence of (partial) overlap of informative voxels between the datasets on DDA while controlling for SNR levels. Critically, we demonstrate that DDA can arise as a result of general differences in SNR between datasets, even when the actual pattern (ground truth) is similar. We argue that since DDA can emerge under these trivial and well-controlled circumstances, authors should be cautious when drawing conclusions based on this effect.

## Materials and methods

In the simulations, we refer to two datasets A and B that both originate from a hypothetical region of interest. Both of these two datasets contain a subset of voxels (the informative voxels) that are differentially responsive to two conditions, and surrounding voxels that show no response to either condition (non-informative voxels). Informativeness, in this context, thus refers to whether a voxel contains relevant information for the condition or not. For each of the datasets, we add varying levels of noise and use MVPA to quantify how this affects both the within-dataset classification accuracy as well as the between-dataset generalizability. In addition, we vary the extent of spatial overlap between the patches of informative voxels between these two datasets to measure the effect of varying overlap and of the number of informative voxels on MVCC. The simulations were run using custom written MATLAB R2014a code (Mathworks Inc, Natick, MA). We use a linear support-vector machine (l-SVM) (Cortes and Vapnik, 1995), based on the LIBSVM algorithm (http://www.csie.ntu.edu.tw/~cjlin/libsvm/) and linear discriminant analysis (LDA) to assess MVCC for these two popular classification algorithms.

### Pattern creation

Both datasets were based on a virtual region-of-interest (ROI) comprising 30 × 30 voxels. Within each dataset a center patch of dimensions 14 × 28 = 392 responsive voxels was created. Half of these voxels showed a response of magnitude 1 to condition 1, and 0.5 for condition 2. The other half of the responsive voxels had the opposite response pattern, with a response magnitude of 0.5 to condition 1 and magnitude 1 to condition 2. This 30 × 30 voxel pattern constitutes the ground truth of a single trial of one condition, see figure 1. In total, 16 trials were created for each condition, totaling 32 trials. This is a low number of trials for machine learning purposes (Varoquaux, 2018), but lies well within the margin of example sizes typically used in MVCC fMRI-studies (Eger et al., 2009; Etzel et al., 2008; Johnson and Johnson, 2014). We then created a trial-specific 30 × 30 Gaussian noise matrix, which was multiplied by a signal-to-noise (SNR) scalar and added to the signal matrix. The SNR scalar is chosen such that the SNR of a dataset follows *SNR*_*dataset*_ = *μ(signal)* / *σ(noise)*, where *μ(signal)* represent the average signal across voxels within the ROI. We varied the SNR levels in both datasets (0 to 0.5, with 0 meaning no signal in the data, i.e. ROI voxels signal set to 0 before adding noise). Note that this SNR represents the signal-to-noise-ratio on a single trial level, and not the temporal SNR that is typically derived from the time course. The overlap of informative voxels between datasets ranges from 0% to 100%, with 100% meaning perfect overlap between the patches informative voxels, thus identical ground truths. By varying these parameters, we simulate several scenarios to test how they affect decoding direction asymmetry.

**Figure 1.**
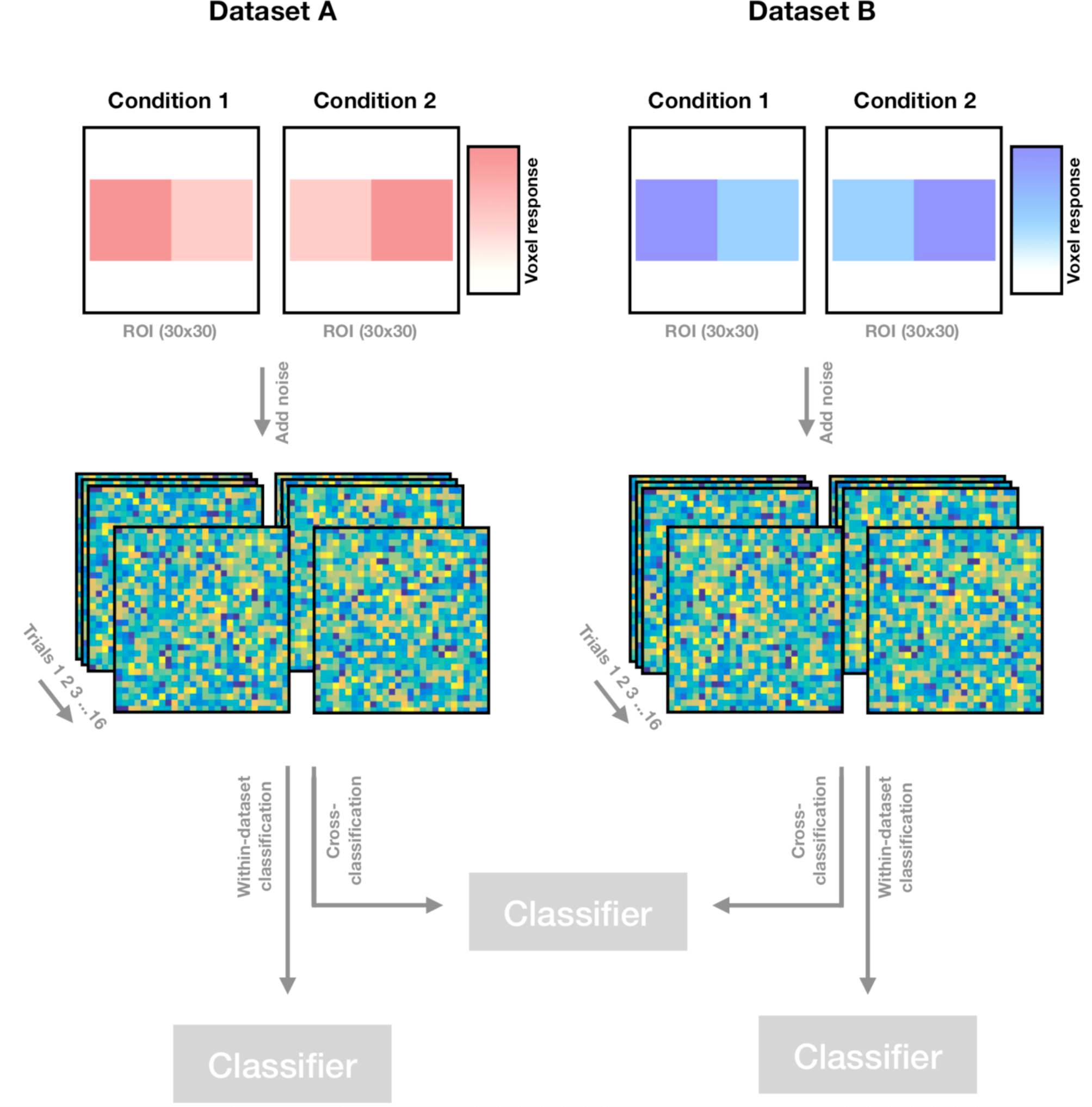
Procedure. Two datasets are created based on a 30×30 ROI. In both datasets, a patch of informative voxels (pink and blue in datasets 1 and 2, respectively) is defined amidst uninformative (i.e. non-responding, white) voxels. Half of the informative voxels show a response of 1 to condition 1 and 0.5 to condition 2, while the other half of the informative voxels shows a response of 0.5 to condition 1 and 1 to condition 2. A Gaussian noise matrix with dimensions 30×30 is created, multiplied by the SNR-scalar and added to the signal patch. This procedure is repeated for each of n trials for each dataset, after which both datasets are Z-scored. Classification is first performed on both datasets independently (within-dataset classification), after which cross-classification is performed on both datasets. See text for details.

### Classification

The 30×30 matrices were transformed in vectors of length 900. The vector of each trial was normalized (z-scored) as to obtain a mean response of 0 and standard deviation of 1 across all voxels within our region-of-interest. We first looked at the within-dataset classification performance. Both a linear SVM as a LDA algorithm were trained and tested using 50 random cross-validations, leaving out 3 trials per condition in each fold. We averaged the performance across folds to obtain the mean prediction accuracy.

Subsequently, we assessed how well one dataset allowed for generalization to the other. We trained the classification algorithm on one dataset as a whole, and predicted the trials of the second dataset through the trained model without cross-validations. Then, we reverse this order by training the classifier on dataset B and testing on dataset A. For each combination of parameter settings that we investigated (noise levels of both datasets, overlap), we repeated the steps mentioned above 20 times (so also generating 20 random pairs of datasets), and we report the average over these repetitions.

### Signal-to-noise variations between datasets

We first examined the effect of different SNRs between datasets on the generalization. We systematically varied the SNR of datasets 1 and 2 while keeping the ground truths in both datasets identical, in steps of 0.01. This way, we could directly investigate in what combination of SNRs we find an asymmetry in decoding direction.

### Overlap between informative voxels

As a second step in our simulations we varied the overlap between the patches of informative voxels in the two datasets, using overlap of 0%, 20%, 40%, 60%, 80% and 100%. In these scenarios, the size of the two patches of informative voxels remains the same, thus leaving the informational content unaltered. As a third and final approach, we investigated the effect of one patch of informative voxels (of dataset B) being a subset of the other patch of informative voxels (of dataset A). We refer to this as dataset partiality, in which we also vary the size of the patch of informative voxels in dataset B, as being 20%, 40%, 60%, 80% or 100% of the size of the first patch. In this scenario, all informative voxels of dataset B correspond to informative voxels of dataset A, but not vice versa. Note that by altering the number of informative voxels in dataset B, the mean signal of this ROI across space is low which affects subsequent SNR computation: following *SNR*_*dataset*_ = *μ(signal)* / *σ(noise)*, the amount of noise that needs to be added to the signal to obtain a given SNR is substantially lower when there are fewer informative voxels in the patch. To make the results comparable between scenarios, the amplitude of noise added to this partial dataset is based on the noise level that would have been used would the dataset have not been partial. The SNR that is reported in the partiality results refers to this noise level approach, not the true SNR of this dataset.

## Results

First, we compared the effect of SNR on DDA in two popular classification algorithms, linear SVM and LDA. We systematically varied the amount of noise in two simulated datasets while cross-classifying between the two, thereby keeping the ground truth of the patterns in both datasets the same. Asymmetry is expressed as the difference in prediction accuracies between training on dataset A and predicting dataset B and vice versa, or schematically: asymmetry = (prediction accuracy dataset A -> dataset B) – (prediction accuracy dataset B -> dataset A). Results are shown in figure 2, where the effect of SNR on DDA is illustrated. The iso-SNR diagonal, which represents the cross-classification performance between sessions with equal SNR (SNR_dataset1_ = SNR_dataset2_), shows no apparent deviation from 0. This indicates that DDA is not simply an effect of a low SNR in both datasets. Rather, DDA arises when the two datasets differ substantially in their signal-to-noise ratios.

**Figure 2.**
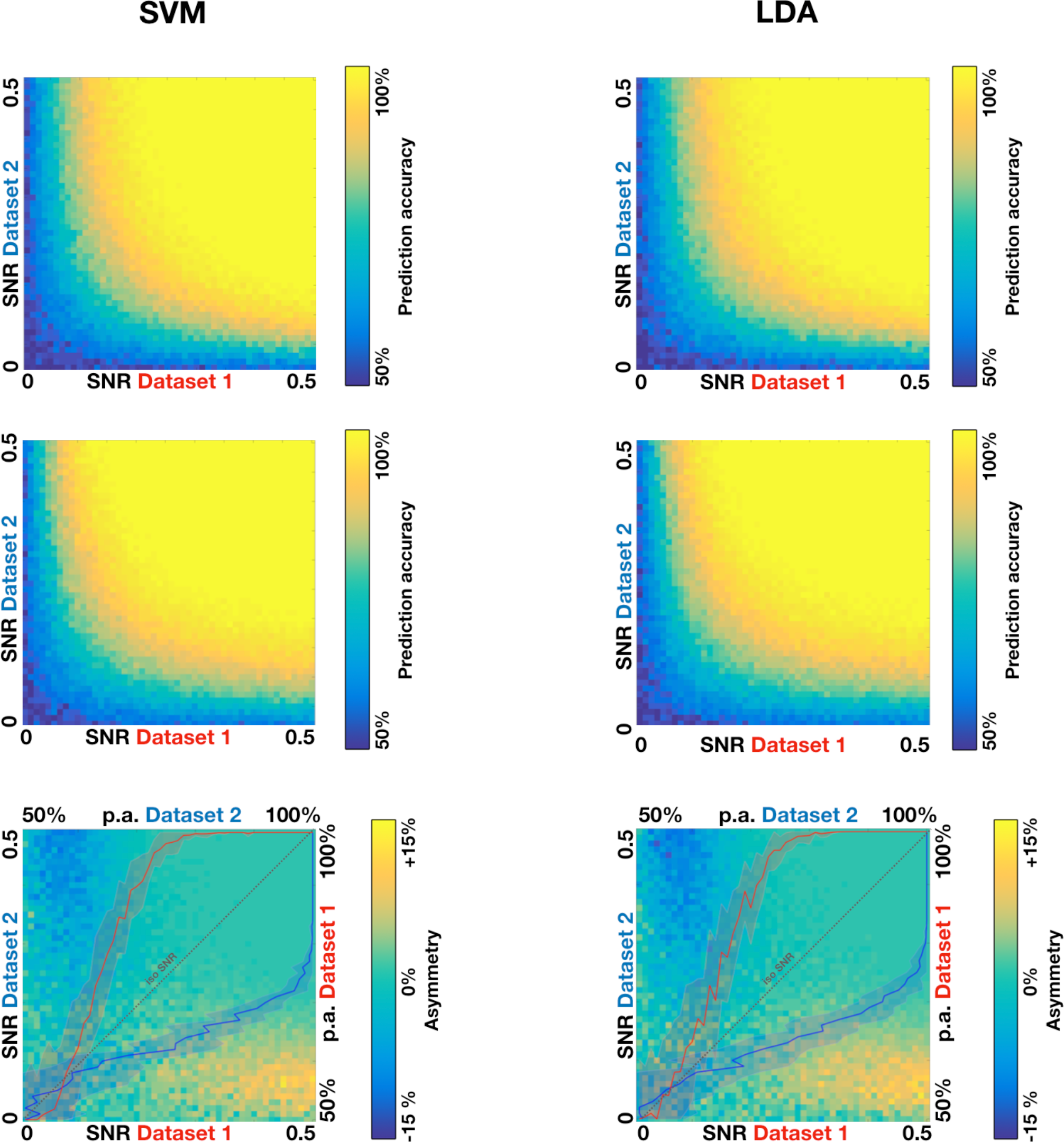
Decoding direction asymmetry as a function of SNR for linear SVM and LDA classification algorithms. The top row panels show the prediction accuracies of training the classifier on dataset A and testing on dataset B, for SVM and LDA classifiers. The x and y axes represent the SNR of dataset A and dataset B, respectively, ranging from 0 to 0.5 in steps of 0.01. The middle row panels show the results of training the classifier on dataset B and testing on dataset B. The bottom panels show the difference (i.e. subtraction) in prediction accuracies between the two generalization directions, thereby directly showing the decoding direction asymmetry. The red and blue line plots show the within-dataset prediction accuracy for dataset A and 2 as a function of SNR (±Standard error of the mean, SEM).

As can be seen, both the linear SVM as LDA classifiers yield decoding direction asymmetry in case of an SNR imbalance between the datasets. In these examples, asymmetry starts to arise when one of the datasets has a low SNR with marginal prediction accuracies, while the other dataset has medium to high trial SNR with high prediction accuracies (>75%). The asymmetry is as such that the generalization is better if classifier training happens on the dataset with low SNR and the cross-validation test is performed on the dataset with high SNR. For example, generalization from dataset A to dataset B is more than 10% better than generalization from dataset B to dataset A when the SNR is around 0.05 in dataset A and above 0.4 in dataset B.

As this scenario has identical ground truths for both datasets, we asked if gradually changing the overlap between the ground truths of both datasets would yield different results. Figure 3 illustrates this effect of varying levels of dataset overlap on DDA. With increasing dataset overlap, generalization asymmetry becomes more pronounced and more skewed towards large SNR differences between datasets. In scenarios with less overlap, the range of SNR values at which DDA arises is broader. In other words, with partial overlap the likelihood of facing DDA at smaller SNR differences increases.

What if the informative voxels of one dataset are a subset of the informative voxels of the other dataset? Figure 4 depicts the effect of this dataset partiality on DDA. When one dataset is comprised of a small number of informative voxels that fully overlap with a larger patch of informative voxels in the second dataset, decoding direction asymmetry arises when generalizing from dataset one to dataset two. Even when the absolute levels of noise in both datasets are equal, a bias in decoding direction can arise as can be seen in the iso-SNR plots of figure 4.

**Figure 3.**
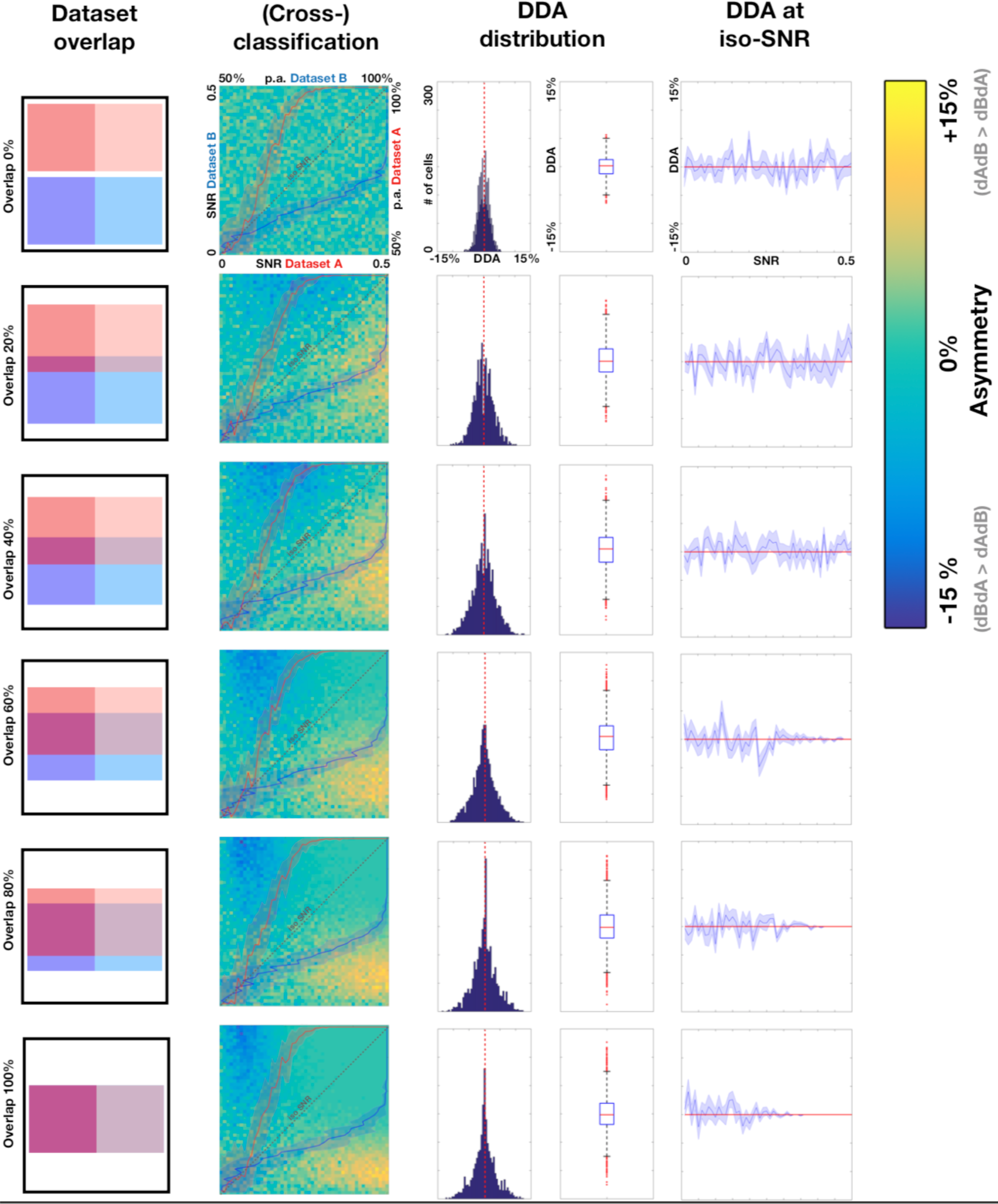
Decoding direction asymmetry as a function of dataset overlap. The left column panels indicate the percentage of overlap of informative voxels between datasets. The cross-decoding matrices show the development of DDA at different SNR levels. To illustrate the distribution of DDA values in the matrix, histograms and boxplots are shown (DDA of 0 excluded for scaling reasons). The rightest plots show the average DDA values at the iso-SNR line ±SEM across the 20 simulations. As can be seen in the cross-decoding matrices, with 0% overlap the cross-decoding and hence DDA yields no systematic results. With increasing overlap, DDA arises but with increasing distance from the iso-SNR line. Also, the magnitude of DDA increases, as can be seen in the DDA distributions. The red line in the histograms and in the iso-SNR DDA plots indicate the DDA = 0 level.

**Figure 4.**
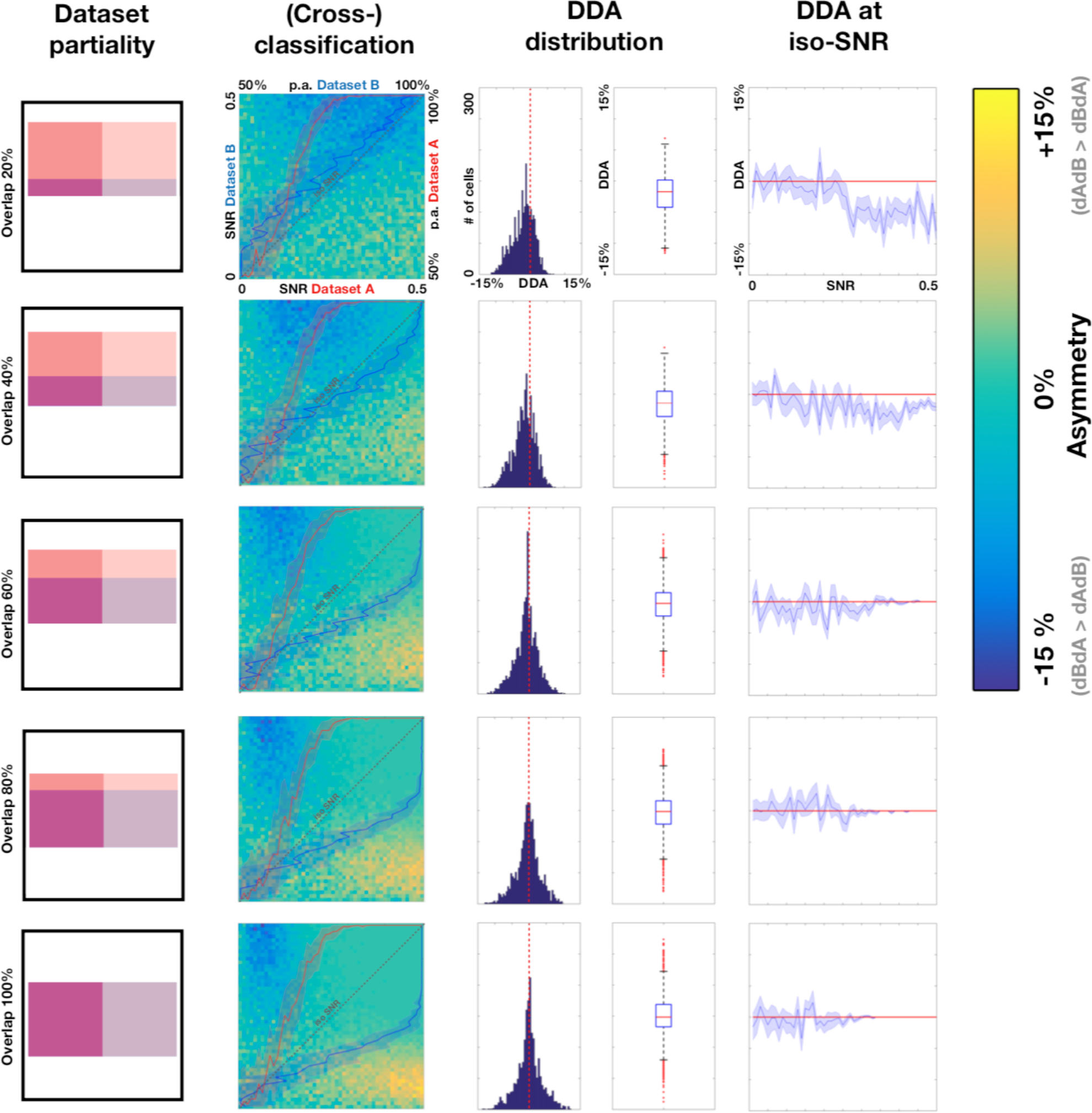
Decoding direction asymmetry as a function of dataset partiality. With dataset B being a small subset of dataset A in terms of spatial distribution, a strong unidirectional decoding direction bias appears in the generalization direction dataset B -> dataset A (blue color). All informative voxels in dataset B contain relevant information about dataset A, but not vice versa. In the 20% overlap scenario, the negative asymmetry bias reaches the iso-SNR line (as can also be seen in the iso-SNR plot), meaning that dataset partiality is a sufficient factor for inducing DDA, irrespective of SNR. For the histograms and boxplots, DDA of 0 is excluded for scaling reasons. The red line in the histograms and in the iso-SNR DDA plots indicate the DDA = 0 level.

## Discussion

In the current paper we investigate the source of decoding direction asymmetry (DDA), a phenomenon that is often reported in studies that perform multivariate cross-classification between neuroimaging datasets. We approached this issue methodologically, first asking to what extent we could reproduce DDA in simple simulations in which we kept the ground truth similar between the two datasets. We showed that differences in signal-to-noise levels between two datasets with equal ground truths is a sufficient condition to yield DDA. This effect is further mediated by the degree of overlap of informative voxels between the two datasets, and by whether information in one dataset is contained in a subset of the informative voxels of the second dataset. In those cases, DDA can arise even when the absolute levels of noise are not very different in the two datasets.

### Interpretation of results

Whereas several neurocognitive or neurophysiological causes of DDA have been proposed (Kaplan et al., 2015), our demonstration that DDA can arise with identical ground truths shows that determining the cause of DDA in a given dataset is not trivial. Finding that dataset A generalizes better to dataset B than vice versa could in theory point to the degree of overlap in informative voxels or partiality of informative voxels in dataset A, which has indeed been proposed in several empirical papers that observed DDA (Eger et al., 2009; Etzel et al., 2008). However, DDA can also be a consequence of SNR differences and thus have no relevance for inferences about overlap in informative voxels or partiality of informative voxels. Therefore, only if one would be able to demonstrate that the noise levels of the two datasets are approximately similar, conclusions that hint towards dataset partiality are warranted. Given that DDA has up to now mostly been observed in situations where there was a clear difference in SNR, researchers should be cautious when attributing DDA to neurocognitive mechanisms.

Similar to the results from our simulations, the cross-classification results that are reported in published work (Akama et al., 2012; Etzel et al., 2008; Johnson and Johnson, 2014; Man et al., 2012; Oosterhof et al., 2012; Quadflieg et al., 2011; van den Hurk et al., 2017) show a stronger performance when the classifier is trained on the noisy or ‘non-preferred’ data (i.e. auditory trials in visual cortex) and tested on the cases from the robust dataset. In our simulations we have shown that a higher SNR yields good within-session classification, but poor generalization to a dataset with lower SNR. Conversely, a within-session classification on a noisy dataset typically results in a lower performance compared to the high SNR data, but the generalization tends to be better. This counterintuitive notion makes sense from a machine learning perspective: classification algorithms such as a soft margin SVM, permit some training examples to be misclassified during the training phase (Mahmoudi et al., 2012). When testing the generalization performance on previously unseen testing examples from the same dataset, the decision boundary is relatively robust to noise in the test set: changing the coefficients of a few support vectors in the training set will likely have little effect on the position of the hyperplane. However, a slight change in orientation of the hyperplane will likely have strongly negative effects on the classification performance on the test set. When this trained model is generalized to a dataset with a higher SNR, however, the decision boundary falls within a wide range of possible boundaries that describe the separation of the classes well. Turning this scenario around, the high SNR dataset has a better within-session classification, but the exact position of the hyperplane is sensitive to a slight adaptation of a few support vectors (Formisano et al., 2008a). This results in a poor generalization to datasets with higher noise levels, even if the ground truth without noise is identical between the datasets.

In the partiality scenarios, we have demonstrated that generalization from the dataset with a small number of informative voxels (dataset A) to the dataset with a large number of informative voxels (dataset B) yields better classification performance than vice versa. As all informative voxels in dataset A contain information about dataset B, a model trained on dataset A will generalize better to dataset B than the other way around.

### Dealing with decoding direction asymmetry

What does this mean in terms of reporting the DDA in neuroimaging studies: are both decoding directions to be averaged, should both outcomes be reported, or is it legitimate to report only the best performance? The key hypothesis that is tested in MVCC is whether the neural representation a brain state is similar in two different contexts (Kaplan et al., 2015). Our findings indicate that the DDA is likely not directly related to experimental or neurobiological factors, but can instead be attributed to aspects of the data itself. More specifically, the lower decoding direction reflects the poor generalization capacity of data that are inherently well separable. We therefore argue that for the sake of transparency both decoding direction results are to be reported, but that the interpretation about the existence of similarity or significant generalization should be based on the best performance direction. This is especially defendable when one can a priori assume that one of the cross-classification directions will outperform the other direction. Thanks to our simulations this decision is no longer based upon unproven theoretical assumptions as was done before (Eger et al., 2009; Etzel et al., 2008), but is grounded in the SNR effects demonstrated in our simulations: generalization will be best when the trained-on dataset has the lowest levels of signal versus noise.

In summary, we demonstrate that decoding direction asymmetry can be a result from an unbalanced SNR between datasets. We argue that since DDA is likely related to factors other than experimental factors of interest, the author is warranted to interpret the strongest decoding direction as evidence for overlapping neural mechanisms at work, especially when the direction of the asymmetry aligns with the direction demonstrated in our simulations.

## Author Contributions

JH and HB conceived and designed the study; JH wrote the simulation code and performed the analyses; JH and HB wrote the manuscript.

